# iResNetDM: interpretable and comprehensive deep learning model for 4 types of DNA modifications prediction

**DOI:** 10.1101/2024.05.19.594892

**Authors:** Zerui Yang, Wei Shao, Yudai Matsuda, Linqi Song

## Abstract

**Motivation:** Despite the development of several computational methods to predict DNA modifications, two main limitations persist in the current methodologies: 1) All existing models are confined to binary predictor which merely determine the presence or absence of DNA modifications, constraining comprehensive analyses of the interrelations among varied modification types. While multi-class classification models for RNA modifications have been developed, a comparable approach for DNA remains a critical need. 2) The majority of previous studies lack adequate explanations of how models make decisions, relying on the extraction and visualization of attention matrices which identified few motifs, and do not provide sufficient insight into the model decision making process.

**Result:** In this study, we introduce iResNetDM, a deep learning model that integrates ResNet and self-attention mechanisms. To the best of our knowledge, iResNetDM is the first model capable of distinguishing between four types of DNA modifications. It not only demonstrates high performance across various DNA modifications but also unveils the potential capabilities of CNN and ResNet in this domain. To augment the interpretability of our model, we implemented the integrated gradients technique, which was pivotal in demystifying the model’s decision-making framework, allowing for the successful identification of multiple motifs. Importantly, our model exhibits remarkable robustness, successfully identifying unique motifs across different modifications. Furthermore, we compared the motifs discovered in various modifications, revealing that some motifs share significant sequence similarities which suggests that these motifs may be subjected to different types of modifications, underscoring their potential importance in gene regulation.

## 1 Introduction

Epigenetic modifications, which encompass mechanisms such as RNA interference (Rosa et al., 2018), histone alterations (Zhang et al., 2021), and the regulation of nucleosome positioning and density (Singh et al., 2021), constitute heritable changes that do not stem from modifications in the DNA sequence itself but rather from the dynamic shifts in DNA accessibility and the architecture of chromatin. Among these, DNA methylation stands out as a critical epigenetic mechanism instrumental in the regulation of gene expression and the safeguarding of genomic stability. Comprehensive research on DNA methylation underscores its indispensability in critical biological processes, including embryonic development, X-chromosome inactivation, and genomic imprinting. Notably, aberrations in methylation patterns are implicated in a spectrum of diseases, with cancer being a prominent example. In malignancies, abnormal hypermethylation can silence tumor suppressor genes (Kim et al., 2019), whereas hypomethylation may activate oncogenes (Tongelen et al., 2017).

DNA modification encompasses a range of biochemical alterations that serve as epigenetic markers, influencing gene expression and cellular function without changing the underlying DNA sequence. Among these modifications, methylation at the sixth position of adenine (6-methyladenine or 6mA) has been observed in prokaryotic and eukaryotic organisms, contributing to the regulation of DNA replication and repair (Wion and Casadesús, 2006). Similarly, 5-methylcytosine (5mC) (Breiling and Lyko, 2015) and its derivative 5-hydroxymethylcytosine (5hmC) (Dahl et al., 2011) are well-recognized in mammalian DNA, playing significant roles in gene silencing and embryonic development. Additionally, 4-methylcytosine (4mC), although less common, has been identified in certain bacteria as part of their unique epigenetic landscapes (Li et al., 2019). Together, these modifications constitute a complex layer of regulatory mechanisms that are essential for the subtle regulation of genetic function and heritable phenotypic variation.

### 1.1 Related Work

The current experimental-based techniques identifying these modifications such as methylated DNA immunoprecipitation (MeDIP) and mass spectrometry can be time-intensive and labor-intensive. In the last decades, several computational methodologies have emerged as alternatives to traditional wet-lab techniques. Most of the machine learning methods are based on support vector machine, Markov model, and decision tree (He et al., 2019; Pian et al., 2020; Pavlovic et al., 2017). As for deep learning, language model and convolution neural network are widely applied to predict 4mC (Xu et al., 2020), 6mA (Zhang et al., 2021) and 5mC (Cheng et al., 2021), respectively. Notably, the advent of the transformer model and self-attention mechanism (Vaswani et al., 2017) has markedly enhanced model performance in predicting DNA modifications (Abbas et al., 2022; Wang et al., 2023; Tsukiyama et al., 2022; Yang et al., 2023).

Despite these advances, most extant models are tailored to recognize only a single type of DNA modification. A notable exception is iDNA-MS (Lv et al., 2020), which stands as the pioneering model employing machine learning to generically predict various DNA modification types. This foundational study delved into the impact of diverse sequential features, such as K-tuple nucleotide frequency component, nucleotide chemical properties, nucleotide frequency, and mono-nucleotide binary encoding, and assessed the efficacy of several machine learning classifiers including Naïve Bayes, Bayes Net, and Decision Tree. Building upon iDNA-MS, subsequent deep learning models like iDNA-ABT, iDNA-ABF, and StableDNAm (Yu et al., 2021; Jin et al., 2022; Zhou et al., 2023) have further honed the precision of DNA modification site detection across multiple species. Yet, these aforementioned models are predominantly binary classifiers, distinguishing only between modified and unmodified nucleotide bases without discerning the specific types of modifications—a multi-classification challenge. In parallel, TransRNAm (Chen et al., 2023) leverages a Transformer combined with a CNN framework to detect twelve distinct RNA modifications, deploying twelve individual classifiers, each dedicated to a single modification type. This approach, while effective for RNA, underscores the absence of a similar, tested methodology in the realm of DNA. Hence, there is a critical demand for a proficient multi-class DNA modification predictor to fill this niche.

In this study, we introduce iResNetDM, which, to the best of our knowledge, is the first deep learning model designed to predict specific types of DNA modifications rather than merely detecting the presence of modifications. iResNetDM integrates a Residual Network (ResNet) (He et al., 2016) with a self-attention mechanism. The incorporation of ResNet blocks facilitates the extraction of local features and empowers the model to achieve a deeper representation of the sequences. Compared to the baseline model, iResNetDM exhibits significant enhancements in performance, achieving high accuracy across all DNA modification types and species. This demonstrates the model’s robust adaptability and efficiency in diverse biological contexts. The main contributions of our work are summrized as below.

1. So far, the existing models can handle at most three DNA modification types simultaneously, namely 4mC, 6mA and 5hmC. Our model takes one step further and adds 5mC based on that.
2. By leveraging the multi-class structure of the model and integrating the integrated gradients (IG) (Sundararajan et al., 2017) technique, we have identified several motifs previously unreported in other studies for the same datasets. This discovery provides deeper insights into the decision-making processes of the model.
3. In our research, we demonstrate that ResNet, although infrequently utilized in this domain, plays a crucial role in enhancing the model’s performance, thereby underscoring its effectiveness and practicality for such tasks. This revelation provides additional options for future studies.

## 2 Materials and method

### 2.1 Datasets selection and preprocessing

The datasets utilized in our study are derived from source (Lv et al., 2020) which comprises 17 datasets across 12 species. For those datasets which are significantly larger than the others, we randomly selected part of them. Given that 6mA is the sole modification occurring on adenine, 6mA-negative dataset was also included to enrich the learning process regarding 6mA. The 5mC dataset for *Z. mays* and *Oryza sativa* Japonica Group cv. Nipponbare is sourced from (Zhang et al., 2020). The combined datasets were input into CD-HIT (Fu et al., 2012) together and those with 80% similarity were removed (Figure 1). To the best of our knowledge, our dataset compilation encompasses all the major known DNA modification types. The details of the datasets can be found in the Supplementary Table S1.

**Figure 1.**
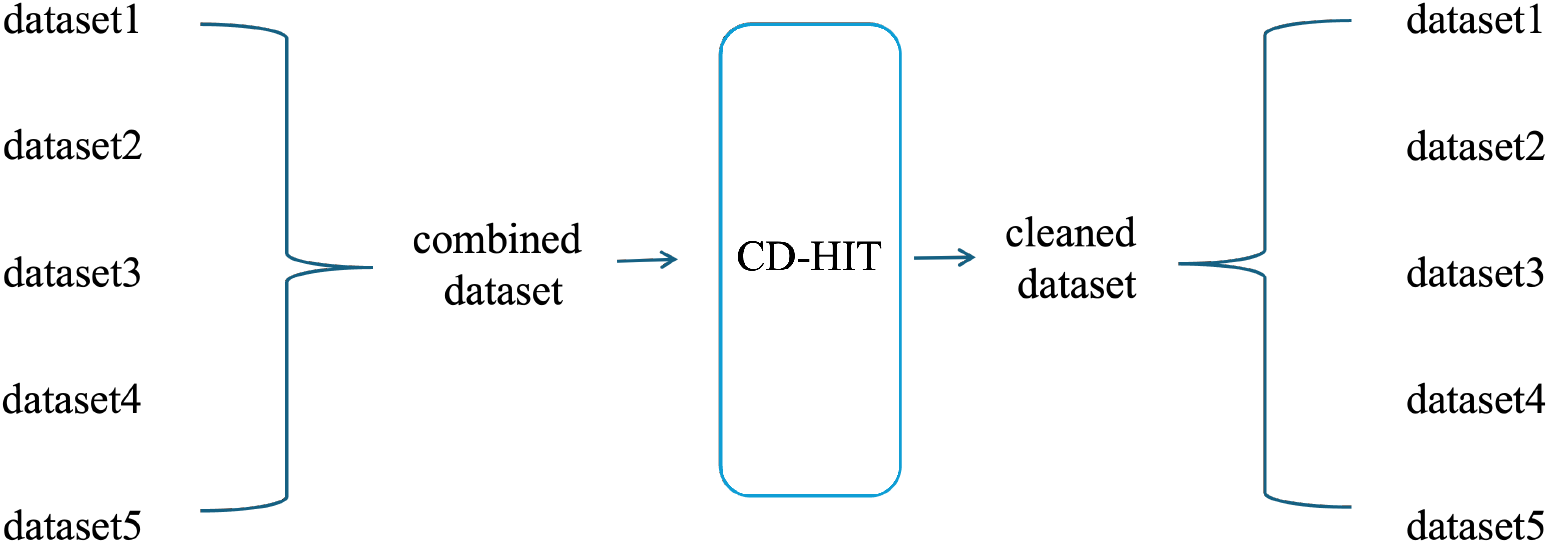
Data pre-processing procedure

All Samples are 41 bp in length with a central modified C or A, flanked by 20 bp regions. Additionally, akin to BERT’s training (Devlin et al., 2019), a [CLS] token is prepended to each sequence before embedding to facilitate subsequent classification.

### 2.2 Model architecture

The architecture of our model (Figure 2)comprises four core components: (1) the Embedding and k-mer Aggregation Layer, (2) the ResNet Layer, (3) the Attention Layer, and finally, (4) the Fully Connected Layer, which synthesizes the output. Initially, each nucleotide in a DNA sequence is converted into a 256-dimensional vector. A k-mer aggregation, which employs a Conv1d layer with kernel size 3, is then utilized to enhance the sequence representation, allowing each nucleotide to capitalize on the neighboring information. Following this, the sequence undergoes a ResNet framework comprising five identical modules that are designed to both extract and synthesize local features, while preserving maximal original data. Subsequently, the output from the ResNet section is processed through a self-attention mechanism, assembling global context. Finally, the DNA sequence’s learned representation is input into a fully connected neural layer, which utilizes a soft-max activation function to execute the prediction.

**Figure 2.**
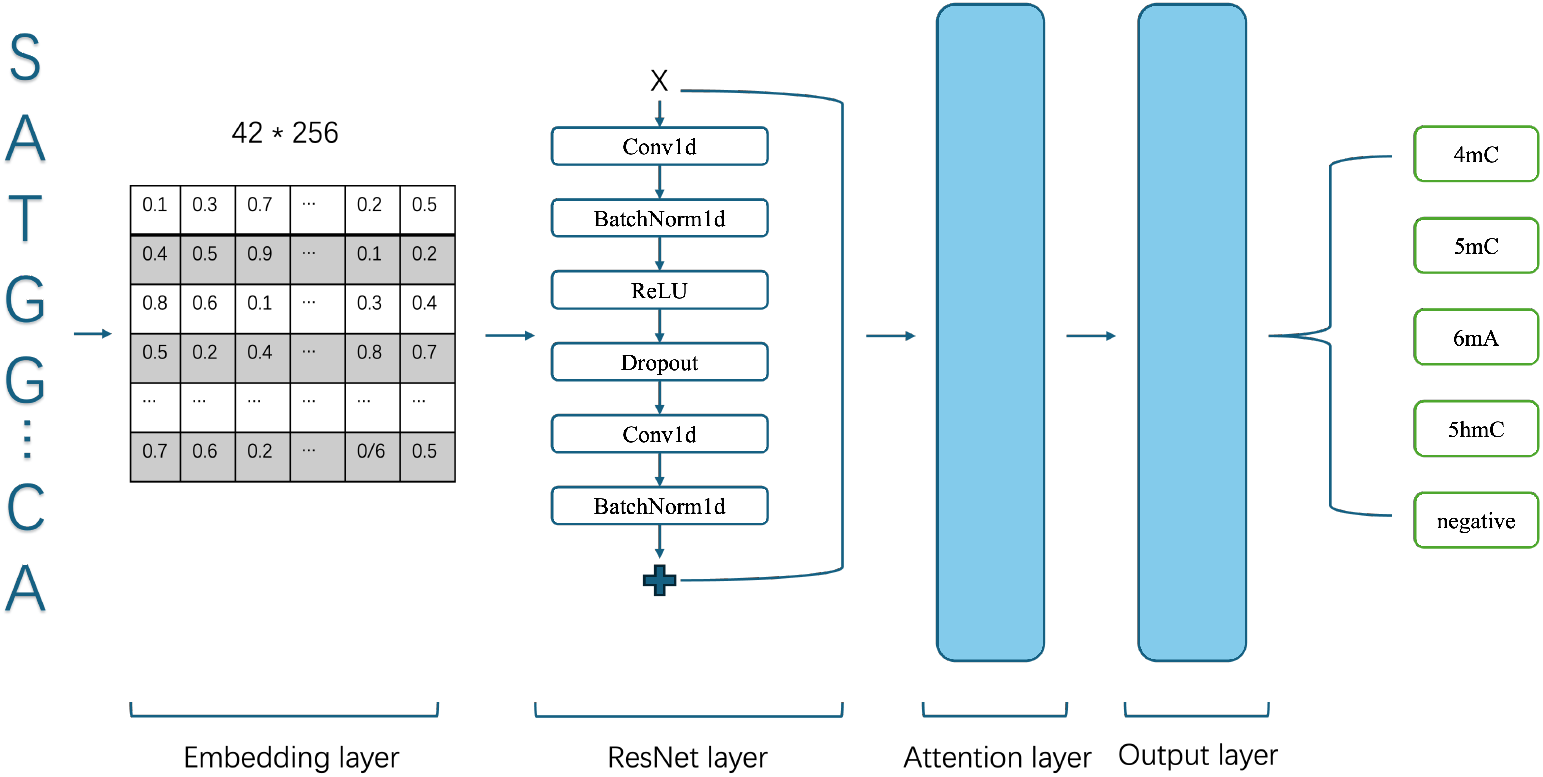
Architecture of iResNetDM. Initially, embedded DNA sequences are fed into a ResNet layer, which is tasked with extracting and learning local features. Subsequently, these features are processed by an attention layer designed to capture global features. Finally, a fully connected layer synthesizes these insights to produce the output.

### 2.3 Residual Networks (ResNet)

Convolutional Neural Networks (CNNs) are extensively utilized in computer vision tasks, offering a powerful tool for capturing complex information from visual inputs. Theoretically, a model with an increased number of CNN layers can encapsulate richer information, leading to enhanced performance. However, this model depth enhancement often encounters the degradation problem; as the architecture deepens, the model may begin to lose track of the original input information, which paradoxically results in performance deterioration after a certain number of layers.

To combat this issue, the Residual Network (ResNet) (He et al., 2016) framework was introduced, utilizing residual connections to bolster the transfer of original information across each network block. Instead of layers learning a direct mapping *H*(*x*), ResNet layers aim to fit a residual function *F* (*x*) conceptualized as *F* (*x*) = *H*(*x*)–*x*. The ResNet layer is thus mathematically articulated as *y* = *F* (*x, W*_*i*_) + *x*.

In instance where *F* (*x*) and *x* have different dimensions, the original ResNet architecture introduces a projection shortcut, resulting in the modified equation *y* = *F* (*x, W*_*i*_) + *W*_*s*_*x* For our purposes, we adapt this framework to maintain the original sequence length of the input data x for straightforward interpretation. We achieve this by incorporating padding in the Conv1d layer, thus preserving the sequence length at 42 throughout the entire computation process.

### 2.4 Multi-head self-attention algorithm

The multi-head self-attention mechanism (Vaswani et al., 2017) initiates the processing of the input sequence by deploying separate and learnable linear transformations. These transformations generate queries (Q), keys (K), and values (V), each uniquely parameterized by their respective weight matrice *W*^*Q*^, *W*^*K*^, *W*^*V*^. Thus, for each attention head indexed by *i*, the transformed representations are expressed as 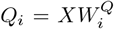, where X denotes the input matrix and h is the total number of attention heads.

Following this, attention scores are computed through the scaled dot product of Q and K, normalized by the square root of the keys’ dimensionality dk, and subsequently passed through a soft-max function to derive the attention weights. This is formally articulated as 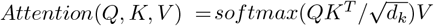

The multi-head attention framework enables parallel processing of these attention computations across the multiple heads, where each head yields an output as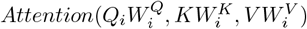. The con-catenated outputs from all heads, *head*_*i*_, are subsequently linearly transformed via the weight matrix *W*^*O*^ to produce the final multi-head self-attention layer output, encapsulated by *MultiHead − Attention*(*Q, K, V*) = *Concat*(*head*_1_, …, *head*_*h*_)*W*^*O*^.

### 2.5 Focal loss

Focal loss (Lin et al., 2017) was employed as the loss function to optimize the model due to the significant imbalance within the dataset, particularly the disproportionately low number of 5-hydroxymethylcytosine (5hmC) samples compared to other categories of DNA modifications. The prevalence of imbalanced datasets can result in a model bias towards the majority class, neglecting the minority class, which is undesirable in medical and bioinformatics applications. Focal loss modifies the cross-entropy loss function by incorporating a modulating factor, which diminishes the contribution of easy-to-classify examples, thereby focusing the model’s attention on the more challenging minority samples. The formula is expressed as *FL*(*p*_*t*_) = *−α*_*t*_ *\** (1 *− p*_*t*_)^*γ*^ *\* log*(*p*_*t*_).

Here, *p*_*t*_ denotes the predicted probability of the actual class *t, α*_*t*_ represents the balancing parameter adjusting the importance of different classes, and *γ* is the focusing parameter that reduces the loss contribution from easy samples. By judiciously selecting *α*_*t*_ and *γ*, focal loss significantly enhances the model’s ability to recognize minority classes, thereby achieving more equitable and accurate predictions in imbalanced datasets.

### 2.6 Integrated gradient

To elucidate the decision-making processes of complex neural networks, particularly in the realm of text data analysis, the IG technique emerges as a powerful tool for attributing predictions to input features. This method offers a transparent and mathematically grounded approach for uncovering the specific contributions of individual input components to the model’s output. By providing a detailed decomposition of the prediction, IG facilitates a deeper understanding of the model’s behavior, enabling researchers and practitioners to pinpoint the exact features that drive the network’s decisions.

Building upon this foundation of interpretability, the application of IG in our experiment involves a systematic approach. First, we identify the neuron corresponding to the input’s label within the output layer, selecting only those samples for which the model assigns a probability larger than 0.3. Subsequently, for baseline establishment, we construct a data point where all feature values are zero. Commencing from the embedding layer, we calculate the gradients of the input data across the entire model by constructing a smooth interpolation path between the baseline and the actual input, comprising 50 interpolation points. For each point along this path, we compute the gradient of the model’s output relative to the interpolated input, accumulating and averaging these gradients to ascertain the integrated gradients for the entire input dataset. Following this, centering on the modification site, segments from each input data and their corresponding integrated gradients are extracted, with flanking regions on both the upstream and downstream sides selected five units in length, resulting in a total flanking region length of ten units. Utilizing UMAP (McInnes et al., 2018) for sequence embedding and DBSCAN (Ester et al., 1996) for clustering all extracted segments, we calculate the mean integrated gradients for each cluster. Finally, we construct the position weight matrix for each cluster to facilitate visualization and employ TOMTOM to compare the sequences identified by our model with motifs discovered by DREME (Bailey, 2011). Segments exhibiting a *p*-value lower than 0.05 are deemed statistically significant.

### 2.7 Nucleotide masking experiment

In the nucleotide masking experiment, each analytical iteration was confined to a singular motif from an individual organism. The token employed for masking purposes was the [CLS] token, which is conventionally appended to the commencement of all sequences as part of the classification process; thus, it does not contribute ancillary information to the model’s predictive deliberations. When masking is implemented within the motifs, the modification sites are prevented from being masked. Conversely, when masking is executed externally to the motifs, the [CLS] tokens, positioned at the onset of each sequence, are judiciously exempted from the masking procedure. To ensure statistical robustness, each experimental iteration was conducted a minimum of ten times to ascertain an average metric for analysis.

## 3. Result

### 3.1 Evaluation

A series of parallel investigations were conducted to ascertain the optimal configuration of ResNet and attention blocks within the network architecture. It was observed that the marginal utility of additional attention blocks diminishes beyond a count of two. Similarly, the incorporation of more than five ResNet blocks was found to inversely impact the model’s efficacy. Consequently, a configuration comprising five ResNet blocks coupled with two attention blocks was uniformly employed throughout our experimental analysis.

The model’s performance was rigorously evaluated using a comprehensive set of metrics: accuracy, precision, recall, F1-score, Matthew’s Correlation Coefficient (MCC), the area under the receiver operating characteristic (ROC) curve, and the area under the precision-recall (PR) curve. The experiment was replicated ten times to ensure reliability, and the mean values were calculated. Collectively, the findings indicate that the model secures an accuracy rate of approximately 0.76. A detailed static of class-specific performance can be found in the Table 1 and Supplementary Figure S1.

**Table 1:**
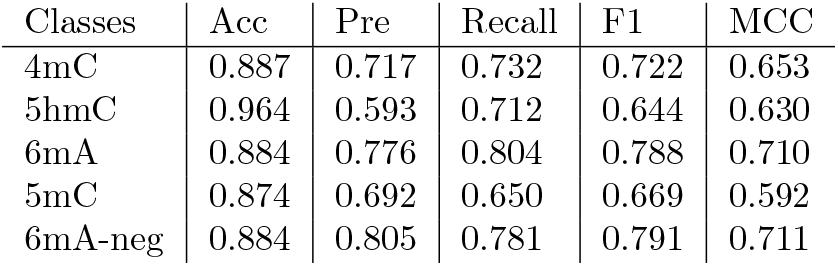
Performance summary of iResNetDM.

| Classes | Acc | Pre | Recall | F1 | MCC |
| --- | --- | --- | --- | --- | --- |
| 4mC | 0.887 | 0.717 | 0.732 | 0.722 | 0.653 |
| 5hmC | 0.964 | 0.593 | 0.712 | 0.644 | 0.630 |
| 6mA | 0.884 | 0.776 | 0.804 | 0.788 | 0.710 |
| 5mC | 0.874 | 0.692 | 0.650 | 0.669 | 0.592 |
| 6mA-neg | 0.884 | 0.805 | 0.781 | 0.791 | 0.711 |

### 3.2 Ablation experiment

To measure the significance of each component within our model, we conducted a series of parallel experiments with various configurations. Initially, we fine-tuned a pre-trained DNABERT (Zhou et al., 2023) model, which served as our baseline, achieving an accuracy of 0.37 (Table 2). This relatively low accuracy underscores the ineffectiveness of self-attention for this task. Inspired by study (Zeng and Liao, 2020), we experimented with incorporating the ResNet architecture. Surprisingly, the integration of ResNet significantly improved model accuracy to 0.763. Furthermore, the addition of each ResNet layer (up to five layers) markedly enhanced model performance, whereas increasing the number of self-attention layers had negligible impact. These findings demonstrate that the ResNet architecture is indispensable for enhancing model functionality in predicting DNA modifications.

**Table 2:** Performance under different configuration.

| Model | Accuracy |
| --- | --- |
| 5 ResNet + 2 self-attention | 0.763 |
| DNABERT | 0.37 |
| 1 ResNet + 2 self-attention | 0.510 |
| 3 ResNet + 2 self-attention | 0.757 |
| 5 ResNet + 5 self-attention | 0.763 |

In addition to evaluating the architectural components of our model, we explored the efficacy of employing focal loss, given its pivotal role in our study. The imbalance in our dataset, particularly the markedly lower amount of 5hmC data compared to other categories, initially led to poor precision and recall metrics when using cross entropy loss, both consistently below 0.1. However, upon implementing focal loss, there was a significant improvement in these metrics—precision increased to 0.593 and recall to 0.712. These results underscore the effectiveness of focal loss in handling class imbalances by focusing more on hard-to-classify instances, thereby substantially enhancing the predictive capabilities of our model in discerning DNA modifications (Supplementary Figure S4).

## 4 Interpretation

The integrated gradient method (He et al., 2017) backtracks from the output to compute the gradient across the model’s architecture. A heightened gradient value signals a more substantial contribution of the corresponding input to the model’s outcome. In our investigation, we harnessed the IG approach to discern potential motifs. To validate our discoveries, we employed an established motif detection algorithm, DREME (Bailey, 2011), as benchmarks. The motif comparison tool TOMTOM (Gupta et al., 2007) was utilized to calculate the *p*-value between the motifs identified by iResNetDM and those found by DREME. Remarkably, the motifs identified by our model align closely with those determined by DREME, demonstrating our model’s capability to recognize motifs as crucial for accurate classification.

Further, the investigation revealed distinct motifs in different species, both in terms of sequence and position (Figure 3 and Supplementary Figure S2). This finding demonstrates that our model is adept at discerning various motifs within the same type of DNA modification, thereby underscoring the model’s robustness and reliability in detecting intricate epigenetic patterns.

**Figure 3.**
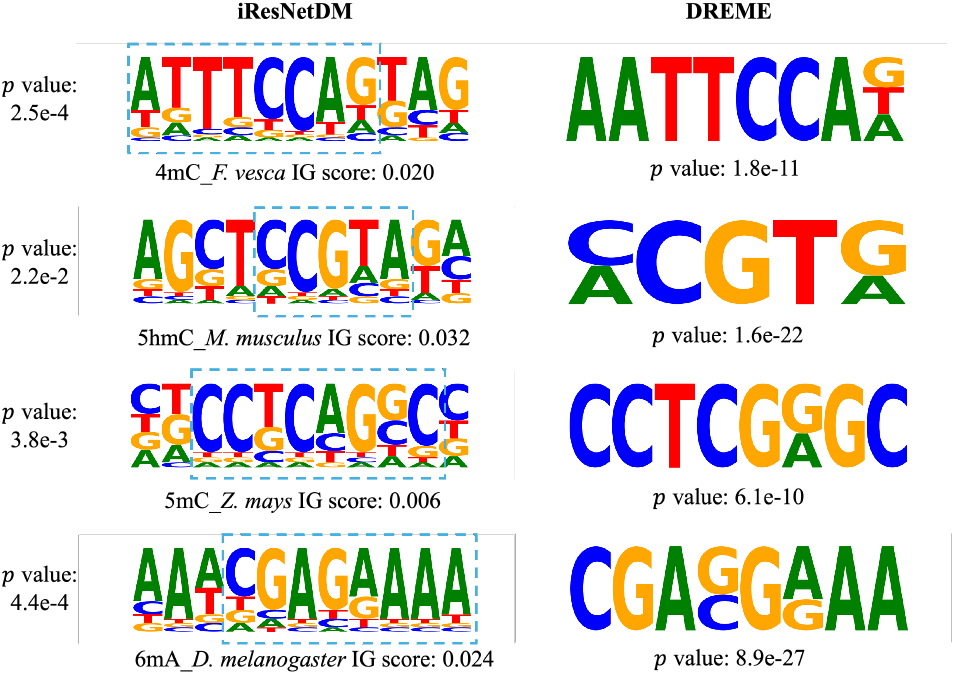
Motif alignment. TOMTOM calculates p-values by comparing observed motifs against a null model comprised of motifs from a target dataset, while DREME uses a one-sided Fisher’s exact test to assess the overrepresentation of motifs within a specific dataset.

To underscore the significance of the identified motifs, we conducted an experimental analysis involving random nucleotide masking both within and external to the detected motifs, subsequently measuring the impact on the model’s recall score. The outcomes unequivocally suggest that nucleotide masking within the motifs exerts a more profound influence on the model’s performance, thereby reinforcing the pivotal role of these motifs (Table 3). Extant research has systematically explored the interrelationships among different species, revealing intricate patterns of genomic interactions (Jin et al., 2021). Building upon these foundational insights, our study employs an integrated model to further elucidate the associations between various DNA modification types. Specifically, for each sequence in our dataset, the output from the last hidden layer was meticulously extracted to compute the average representation for each class of DNA modifications.

**Table 3:** Impact of Nucleotide Masking on Model Recall Scores by Motif and Species. The nucleotides marked in red stands for the modification sites.

| Modification | Species | Motif | Mask inside(std) | Mask outside(std) |
| --- | --- | --- | --- | --- |
| 4mC | *F. vesca* | ATTTCCAG | 0.694 (0.065) | 0.706 (0.040) |
| 5hmC | *M. musculus* | CCGTA | 0.229 (0.031) | 0.403 (0.050) |
| 5mC | *Z. mays* | CCTCAGGC | 0.691 (0.049) | 0.787 (0.066) |
| 6mA | *D. melanogaster* | CGAGAAAA | 0.891 (0.031) | 0.920 (0.028) |

The correlation heatmap (Figure 4) presented underscores a consistent negative correlation between 6mA and other DNA modifications, suggesting an antagonistic regulatory mechanism within the genomic landscape. Conversely, modifications occurring on cytosine—namely 5mC, 5hmC, and 4mC—demonstrated strong mutual correlations. This observation is indicative of co-occurrence and possibly cooperative interactions in certain genomic regions that are extensively modified by multiple types of DNA modifications. To substantiate this hypothesis, our analysis extended to the examination of motifs identified by the DREME algorithm (Supplementary Figure S3). This inquiry uncovered several motifs that are common across 5mC, 5hmC, and 4mC, thereby underscoring their potential significance in epigenetic regulation. These motifs, prevalent among highly correlated modifications, may serve as crucial sites for the orchestration of epigenetic marks that modulate gene expression in a concerted manner. Our findings suggest that these regions might play pivotal roles in cellular processes, warranting further investigation into their biological implications and their overarching impact on gene regulation dynamics.

**Figure 4.**
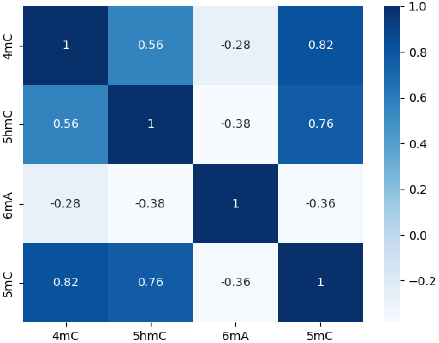
Correlation between different modifications revealed by iResNetDM

To validate the functionality and efficiency of each component within our model, we extracted the outputs from each section and employed t-SNE (Laurens and Hinton, 2008) for dimensionality reduction and visualization (Figure 5).The sequential analysis of outputs from various layers of the neural network delineates the integrated functionality and complementary nature of the ResNet and self-attention mechanisms in processing complex features. The embedding layer, serving as the initial feature representation, exhibits a dispersed clustering of DNA modifications, setting a baseline for subsequent feature enhancement. This layer underscores the raw state of the data, indicating the necessity for substantial transformations to achieve meaningful discriminability.

**Figure 5.**
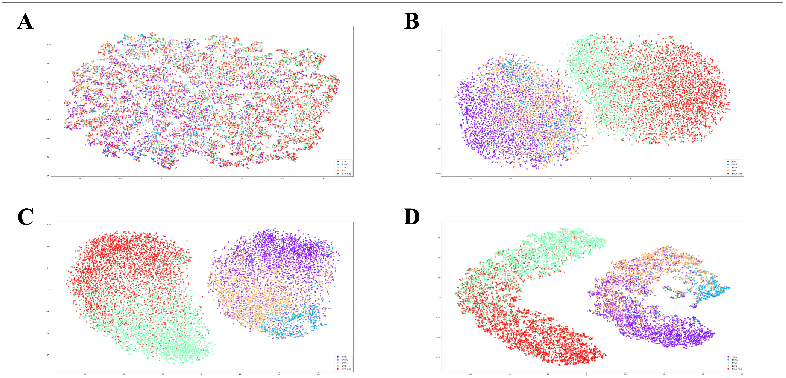
t-SNE visualization of the output extracted from (**A**): the embedding layer (**B**): the ResNet layer (**C**): the attention layer (**D**): the output layer.

As the first step in this transformation process, the ResNet layers initiate a basic yet crucial separation of features. Although this separation marks an improvement over the embedding layer, it remains insufficient for high discriminability, reflecting the intrinsic complexity of the data involved. The ResNet layers thus establish a fundamental framework for feature differentiation but do not fully capitalize on the potential for fine-grained classification.

Building upon the groundwork laid by the ResNet layers, the attention layers significantly refine the feature representations. By effectively concentrating on the most influential features, these layers enhance the clarity and distinction of the clusters, achieving superior clustering performance.

The apex of this process is observed in the output from the last hidden layer, where the integration of diverse features results in distinct and refined clustering. This output not only signifies the network’s capacity to synthesize and enhance features progressively but also highlights its ability to achieve highly effective classification capabilities.

## 5 Discussion

In the current DNA modification research, existing models are primarily built upon binary classification tasks. While these models are adept at distinguishing the presence or absence of DNA modifications, they fall short in analyzing the intricate relationships among different types of modifications, thus limiting the scope for more comprehensive analyses. To address this limitation, we introduced the iResNetDM model. This model is not only capable of distinguishing between various classes of DNA modifications but also leverages its integrated features to perform a holistic analysis of the relationships among these modifications. Furthermore, the utilization of IG techniques enhances the model’s interpretability, thereby enriching our understanding of the underlying biological processes.

In this investigation, we have devised an innovative model that demonstrates exceptional efficacy in discerning four prevalent DNA modifications. This study underscores the utility of the ResNet architecture in predicting DNA modifications and marks a novel application of ResNet in this domain. The conventional self-attention mechanism, which typically shows promise in other contexts, faltered in our study. However, the iResNetDM model, with its impressive accuracy across various organisms, successfully identified several motifs, thereby highlighting its robustness and broad applicability.

This work pioneers the use of IG techniques in discerning potential motifs associated with DNA modifications, aligning with similar research endeavors to a degree (Song et al., 2021). Notably, our findings exceed in both the breadth and the quality of the identified motifs compared to existing studies. For instance, certain research (Jin et al., 2022) lacks the expansive data utilization that characterizes our approach, potentially limiting statistical robustness. Leveraging the insights gained from systematic explorations of genomic interactions across different species (Jin et al., 2021), our study has applied an integrated model to elucidate the complex relationships among various DNA modification types. By extracting the output from the last hidden layer for each sequence, we have computed detailed representations that have enabled us to highlight specific patterns of modifications.

Our analysis has revealed a distinct pattern of interaction among modification types. Specifically, a consistent negative correlation between 6mA and other DNA modifications suggests an antagonistic regulatory mechanism, while strong mutual correlations among cytosine-based modifications (5mC, 5hmC, and 4mC) indicate potential cooperative interactions within certain genomic regions. These regions, extensively modified by multiple types of DNA modifications, were further analyzed to identify common motifs through the DREME algorithm. The identified motifs underline the potential significance of these modifications in epigenetic regulation, suggesting that they may serve as crucial sites for orchestrating epigenetic marks that modulate gene expression.

Such findings not only underscore the utility of integrated gradient techniques in enhancing our understanding of DNA modifications but also suggest pivotal roles for these regions in cellular processes, warranting further investigation into their biological implications and their impact on gene regulation dynamics.

Despite this progress, constraints linked to the available datasets curtailed the scope of motif identification for other DNA modifications, such as 6mA, while some, like 5fC and 5CaC (Barnett et al., 2014), were omitted due to the absence of corresponding datasets. Anticipation abounds for the application of our model to novel datasets related to diverse DNA modifications, particularly those associated with adenine, such as 6hmA (Xiong et al., 2019), to further substantiate its performance.

## 6 Conclusion

In this work, we have introduced iResNetDM, a deep learning model tailored to predict and distinguish between four specific types of DNA modifications. Our results underscore the capability of iResNetDM to deliver high-performance metrics, demonstrating robust adaptability and efficiency across various DNA modifications and species. This study represents a significant step forward in the field of bioinformatics, particularly in the understanding and prediction of complex epigenetic modifications.

Key contributions of this research include the development of a model that can handle multi-class DNA modification predictions—a first in the field. By integrating ResNet and self-attention mechanisms, iResNetDM has shown not only to enhance prediction accuracy but also to provide deeper insights into the decision-making processes underlying DNA modification. This was achieved through the innovative application of the IG technique, which facilitated a clear understanding of the contributions of individual sequence components to the predictions made by our model.

## 7 Data and Code Availability

The datasets and code of our work will be released on GitHub at https://github.com/yaoge777/iResNetDM following publication. For the convenience of future studies, the curated dataset will also be released on GitHub.

## 8 Contribution

Zerui Yang initialized the project, implemented the model, and conducted the experiments. Linqi Song and Yudai Matsuda supervised the project. Linqi Song, Yudai Matsuda and Wei Shao critically reviewed the manuscript.

