## Supplementary material for "iResNetDM: interpretable and comprehensive deep learning model for 4 types of DNA modifications prediction": SI: SI.docx

| 4mC | *C. equisetifolia* | 319 | 46 |
| --- | --- | --- | --- |
| 4mC | *F. vesca* | 8750 | 1250 |
| 4mC | *S. cerevisiae* | 1723 | 247 |
| 4mC | *Tolypocladium* | 8750 | 1250 |
| 4mC-total |  | 19542 | 2793 |
| 5hmC | *H. sapiens* | 2053 | 291 |
| 5hmC | *M. musculus* | 3191 | 456 |
| 5hmC-total |  | 5244 | 747 |
| 5mC | *Z. mays* | 8750 | 1250 |
| 5mC | NIP | 8750 | 1250 |
| 5mC-total |  | 17500 | 2500 |
| 6mA | *A. thaliana* | 2625 | 375 |
| 6mA | *C. elegans* | 2625 | 375 |
| 6mA | *C. equisetifolia* | 2625 | 375 |
| 6mA | *D. melanogaster* | 2625 | 375 |
| 6mA | *F. vesca* | 2446 | 350 |
| 6mA | *H. sapiens* | 2625 | 375 |
| 6mA | *R. chinensis* | 512 | 375 |
| 6mA | *S. cerevisiae* | 2625 | 375 |
| 6mA | *T. thermophile* | 2625 | 375 |
| 6mA | *Tolypocladium* | 2625 | 375 |
| 6mA | *Xoc BLS256* | 2625 | 375 |
| 6mA-total |  | 26808 | 4100 |
| 6mA-neg | *A. thaliana* | 2625 | 375 |
| 6mA-neg | *C. elegans* | 2625 | 375 |
| 6mA-neg | *C. equisetifolia* | 2625 | 375 |
| 6mA-neg | *D. melanogaster* | 2625 | 375 |
| 6mA-neg | *F. vesca* | 2446 | 350 |
| 6mA-neg | *H. sapiens* | 2625 | 375 |
| 6mA-neg | *R. chinensis* | 512 | 375 |
| 6mA-neg | *S. cerevisiae* | 2625 | 375 |
| 6mA-neg | *T. thermophile* | 2625 | 375 |
| 6mA-neg | *Tolypocladium* | 2625 | 375 |
| 6mA-neg | *Xoc BLS256* | 2625 | 375 |
| 6mA-neg-total |  | 26808 | 4100 |
| Total |  | 95902 | 14240 |

Table S1: Statistic of datasets

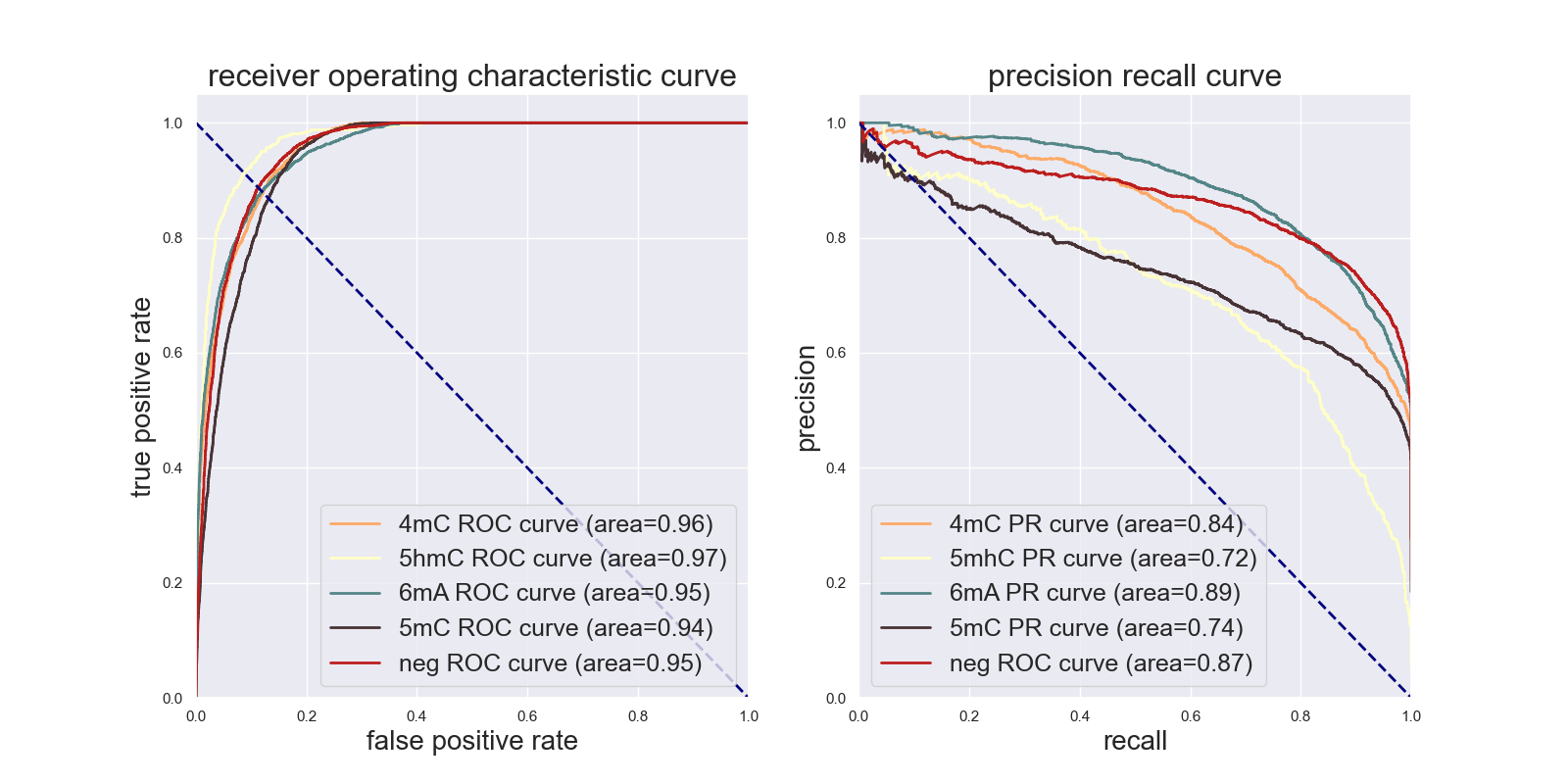

**Figure S1**: Receiver Operating Characteristic (ROC) and Precision-Recall (PR) Curves for the Model

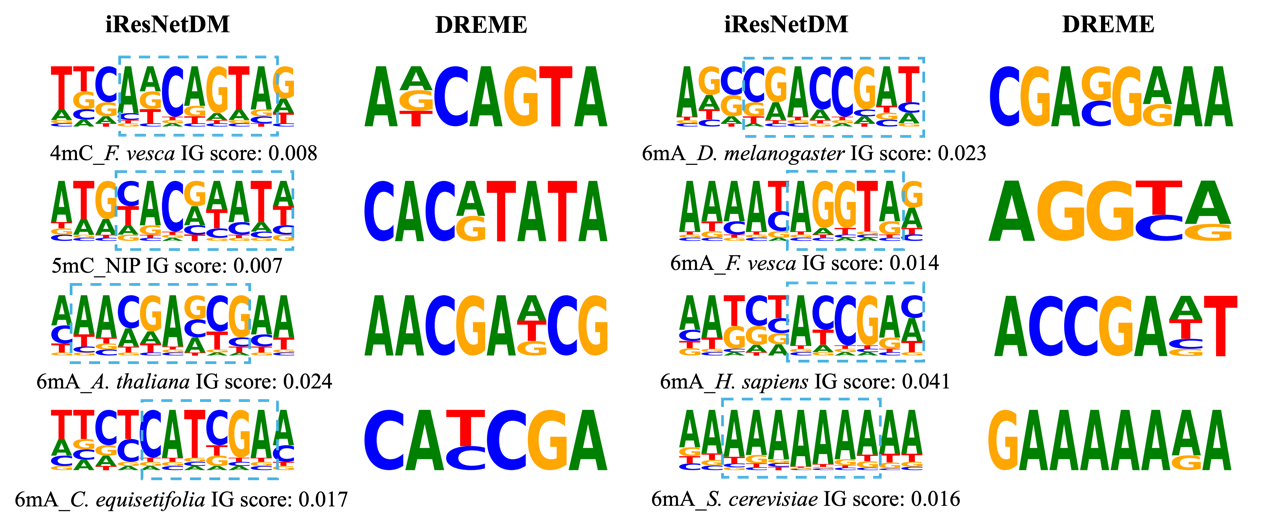

**Figure S2**: Motif alignment. only motifs found by DREME with p-value lower than 0.05 and motif pairs with p-value computed by TOMTOM lower than 0.05 are selected

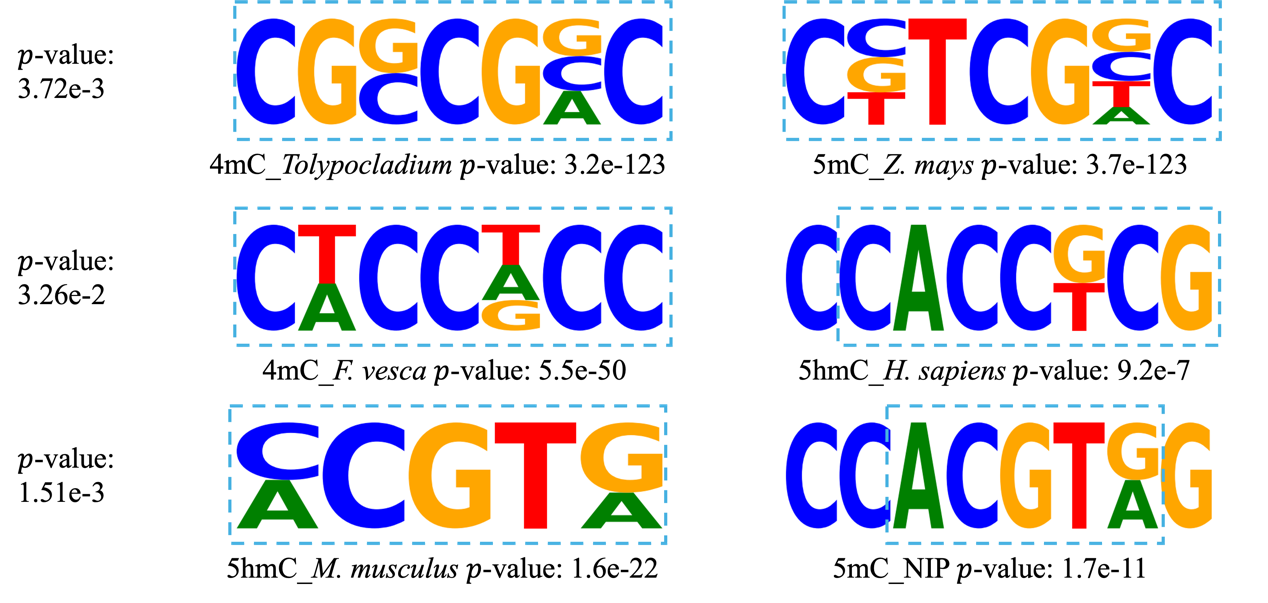

**Figure S3**: Alignment between some motifs of 4mC, 5mC, 5hmC

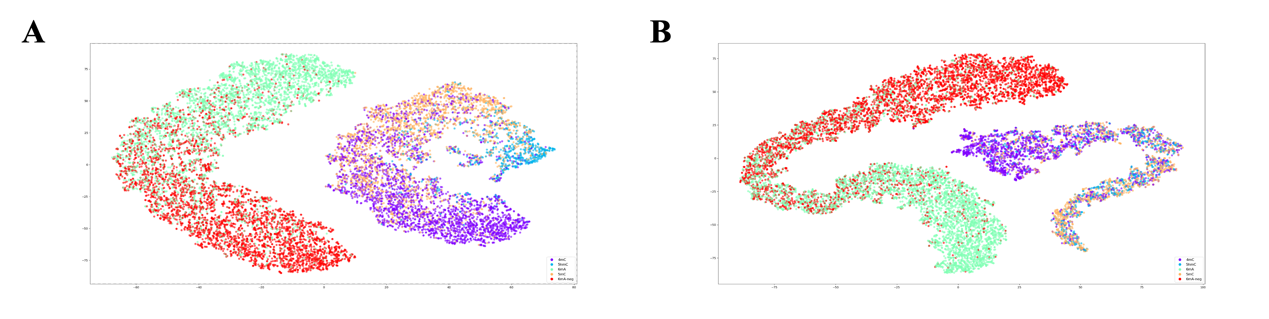

**Figure S4**: t-SNE visualization of the outputs from the model's output layer, comparing the effectiveness of different loss functions: (**A**) focal loss and (**B**) cross-entropy loss. The visualization illustrates that focal loss markedly enhances the model's ability to distinguish 5-hydroxymethylcytosine (5hmC), particularly in cases where sample sizes are limited
